# Feeding behaviour, species associations and natural diets of 10 Cyprinid fish species from South-West Sri Lanka

**DOI:** 10.1101/2023.07.12.548677

**Authors:** Koenraad Kortmulder, Peter van der Wiele

## Abstract

Sri Lanka has been a favored region for studies of fish assemblages in tropical streams. Food, habitats, reproduction and species segregation have been in focus. The present paper introduces a new approach, comparing feeding behavior of 10 indigenous cyprinid species as observed in aquaria under standardized conditions. It is found that species with similar feeding habits (as to substrate preferences and some characteristics of snapping and processing) are largely separated in their typical habitats in the wild. This conclusion is similar to what others, working with largely the same species in the same region, have found, namely that co-existing species differ in their food preferences. But our conclusion was arrived at independently on the basis of different data. Advantages of behavior studies in this context are discussed, such as identification of evolutionary adaptations in behavior and added information on *where* in their common habitat co-occurring species collect certain food items that are commonly found in gut content analyses.

## Introduction

Studies on fish assemblages in tropical streams are many, and Sri Lanka, particularly the SW wet zone of the island, has been a favored region for them. The main thrust of this research has been to explain the richness of the fish faunas in term of species segregation. Most studies focus on diets, which are read from intestinal contents of specimens caught in the wild. Moyle & Senanayake (1984), however, also investigated a wide range of aspects, including morphological features such as have to do with food choice, and diverse characteristics of the immediate aquatic environment: water depth and velocity, and substrates. De Silva *et al*. (1977 and 1980) described gut lengths and gill rakers of four indigenous Cyprinids and noticed comparatively prominent esophageal teeth in *Puntius dorsalis*. Schut *et al*. (1983) described typical habitats of eight Cyprinid species, and calculated degrees of co-occurrence. They distinguished three or four species associations, ranging from hill marshes at relatively high altitudes to lower stream stretches and lowland marshes around flood level. A less complete analysis of lowland species is offered by Kortmulder & Robbers (2011). Weliange and Amarasinghe (2003) studied daily rhythms of feeding in a lacustrine environment, and demonstrated that ignoring them unduly influences the calculation of overlap between species. Only a few studies supplemented analyses of intestinal contents with samples of the environment taken along with the collection of the fishes, in order to investigate *preferences* on the part of the fishes (Shirantha *et al*. 2005; Vijverberg *et al*., 2019). Vijverberg *et al*. (2017) also made direct observations of foraging fish in the field. A somewhat neglected paper by Fernando (1956) reports, besides data on gut contents, aquarium experiments and qualitative but expert surveys of potential food organisms at all collection locations.

Without these refinements, the diets of at least some species have proven to be very variable from one location to another and/or in different seasons. For instance, *Pethia nigrofasciata* guts from respectively Kalu and Kelani River systems contained 6 and 24% volume diatoms and 26 resp. 14% higher aquatic plant material (Shirantha *et al*. 2005). In the same species, Moyle and Senanayake (1984) found less than 1% diatoms, and De Silva & Kortmulder (1977) 75% diatoms and 25% green algae. The same three papers report 4% or less animal matter, but Vijverberg *et al*. 2017 found up to 25%. Similar discrepancies could be offered for *Puntius dorsalis* (Vijverberg *et al*. 2019; Moyle & Senanayake, 1984; De Silva *et al*. 1980; Fernando, 1956).

One explanation might be that segregation of these species is not by selection on food, but for instance by predation or by availability of favorable spawning places. In that case the full-grown fishes could just be opportunists: eating what is nearest to their mouths, and yet be selected for being different. It is important to note, however, that Shirantha *et al*. (2005) found more agreement between *Pethia nigrofasciata* gut contents from Kalu and Kelani rivers when the environmental samples were brought into play. (Comparison with Vijverberg *et al*. 2019 is not possible because food categories were different). Moreover, if selection has not been for food, how could morphological features such as highly developed gill rakers be explained? Thus, the assumption that food is an important segregating factor, is quite reasonable. However, for future research, the combined collection of fish and environmental samples is to be recommended, in addition to observations of the fish as to the substrates from which they feed, before they are disturbed by the catching procedures.

A new approach is presented in this paper, concerning laboratory experiments and focusing on the species-specific feeding behaviors. The presence in our lab -in the early eighties -of 10 out of the 12 ‘barb’ species then known^2^ to occur in SW Sri Lanka: *Puntius kamalika, Puntius chola, Puntius bimaculatus, Puntius dorsalis, Puntius vittatus, Puntius titteya, Pethia nigrofasciata, Pethia reval, Systomus pleurotaenia* and *Dawkinsia singhala*, inspired precise investigations in aquaria. All species were observed under similar conditions and according to a fixed scheme and provided with the same food^3^. This principle brought out the typical properties of foraging behavior of each species, which turned out to be highly specific.

## Material and methods

Living specimens of *Puntius bimaculatus, P. dorsalis, P. kamalika, P. chola* and *Systomus pleurotaenia* used for this study had been imported directly from Sri Lanka about one year earlier. The other species used: *Puntius titteya, P. vittatus* ^4^, *Pethia nigrofasciata, Pethia reval* ^5^ and *Dawkinsia singhala* represented long established lines in the lab. Behavior of all kinds was impressively conservative in our lab stocks, even over generations.

The observations on which the present study is based, were all made in aquaria of 125 × 40 × 50 cm (lxbxh) with glass on all sides. Stocks were kept in similar tanks in the same thermo-regulated room (*c*. 24°C). Artificial light was provided 08 am -22 pm. In part I of the program, groups were observed; in part II a single individual, a male or a female (series II♂ and II♀ respectively). A *series* comprised all 10 species. Series were repeated in such a way that all species were observed under all conditions of bottom material. In the groups, one individual was followed. When it proved non-representative, for instance when it was territorial in a non-territorial crowd or when it was exceptionally inactive as compared to its companions, another one was chosen instead. When all fish of a group were relatively inactive at feeding, observation time could be extended beyond its standard length in order to get sufficient numbers of snaps. ‘Inactive’ means less than 2 or 3 snaps in the allotted time. For statistical purposes, we needed at least 20 snaps in the 10 observation periods of a series (see below, p. 00). Normal observation periods were constant within each series, but could vary between series from 5 to 6 or 7 minutes. Snapping type characteristics (depth, accompanying movements) and food processing after the snap (chewing, spitting, sieving) were scored in separate periods during each session. A complete session lasted 30 minutes (or less, depending on the series) and was begun with some minutes rest, to allow the fish to quiet down after the arrival of the observer(s). In the next section (Results), a *sample* means the sum over a whole series.

Observations were made, between 9.15 and 16.00 hrs, on ten consecutive days, on all 10 species in a fixed order, but beginning with the next species in the row each day, so that after 10 days (one series) all species had been observed at all times of the working day. The fish were fed at 9.00 am (sparingly) and 16.00 pm with commercial dry food or live *tubifex*. During part II of the study (single individuals) only one kind of dry and no live food was offered. Observations were made January through August 1982 (part I) and May through November 1983 (II). Pauses of at least a few days separated series which involved reorganization of the tanks or moving of the fish to another tank.

The observation tanks were provided with bottom layers of coarse sand, ‘gravel’ (very small grains, slightly larger that those of sand and rounded) or sand-colored PVC tiles of 5 × 5 cm each. The first two were applied covering the whole bottom (series Ia and Ib), or all three were combined two by two on bottom halves (final two series of part I; part II; substrates always swapped in consecutive series). All tanks had living plants, one or more of *Vallisneria asiatica, Hygrophila verna*, low *Cryptocoryne, Microsorum pteropus* and, at least, filamentous algae^6^.

Feeding behavior occurred in episodes separated by other activities and consisted of snaps, performed singly or in sequences, the latter at the same place as the first snap or describing a trajectory over the substrate. In between snaps, either within or between sequences, a specific searching posture could be performed, with close fixation of the substrate and when at the bottom a 30^°^ head-down orientation, or a normal swimming posture could be maintained. As a measure for the preponderance of the searching posture, we scored the percentage of all bottom feeding episodes without snaps that were performed in this posture.

## Results

### 1. Substrate choice

We distinguished six feeding substrates: bottom, plant, wall (any vertical surface except plants), water column, water surface, and plant-at-water-surface. The first three together constitute the ‘fixed substrates’; the other three are referred to as ‘free’. Twenty samples were available for each species. Samples containing less than 20 snaps were excluded from the statistical treatment (table 1). The table presents, for each species, the percentages ‘bottom’ and ‘bottom+plant+wall’.

**Table 1.**
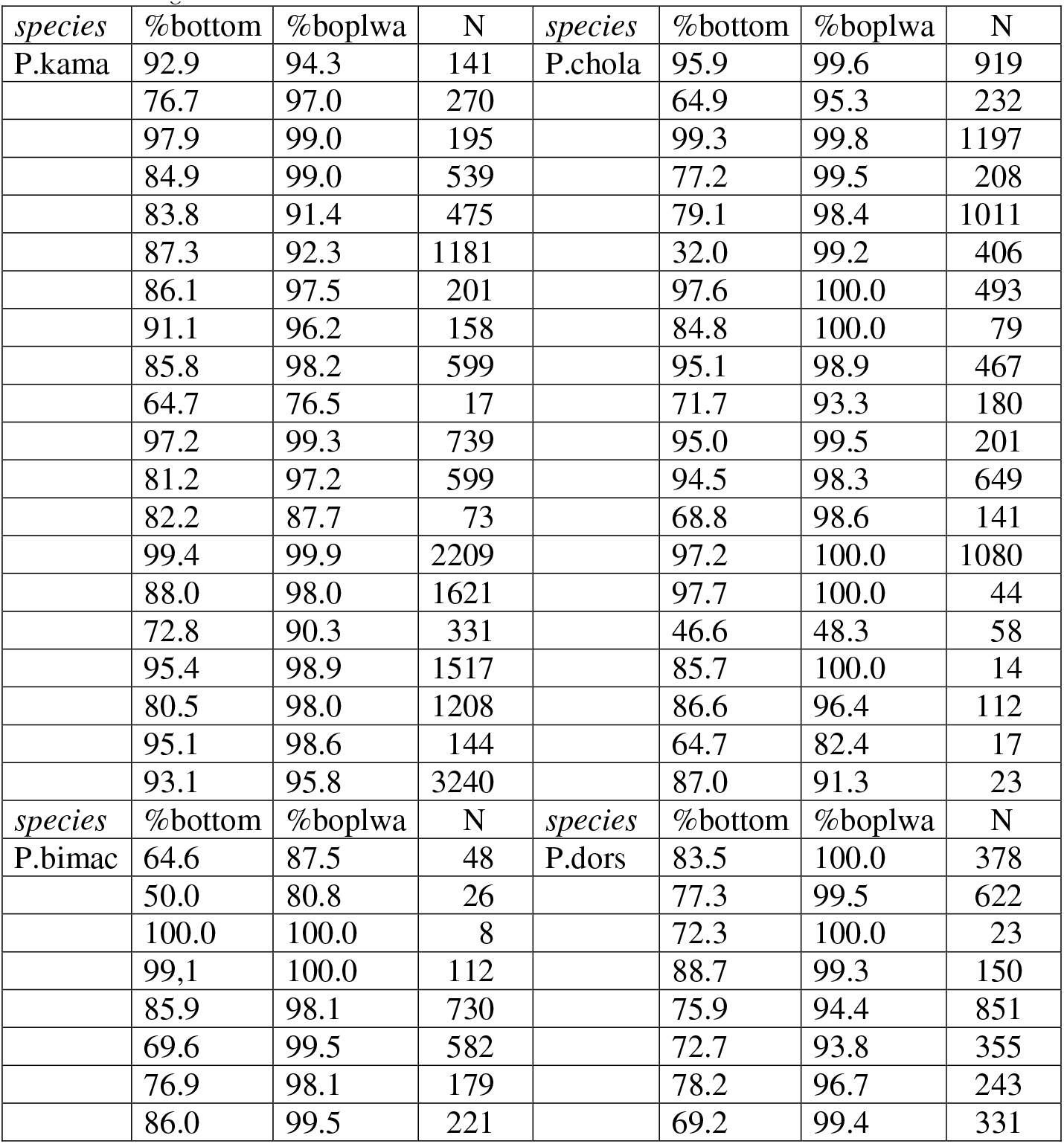
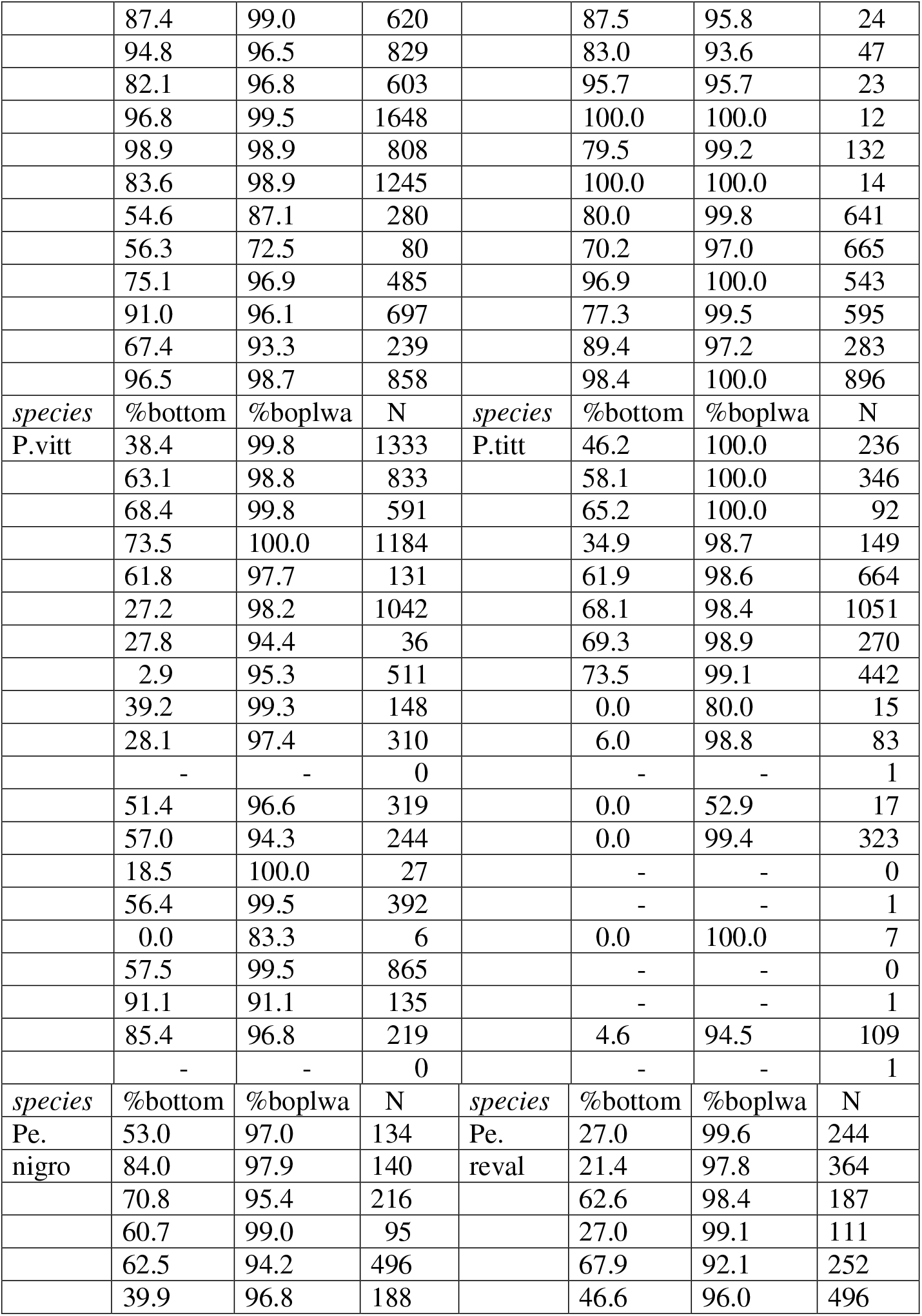
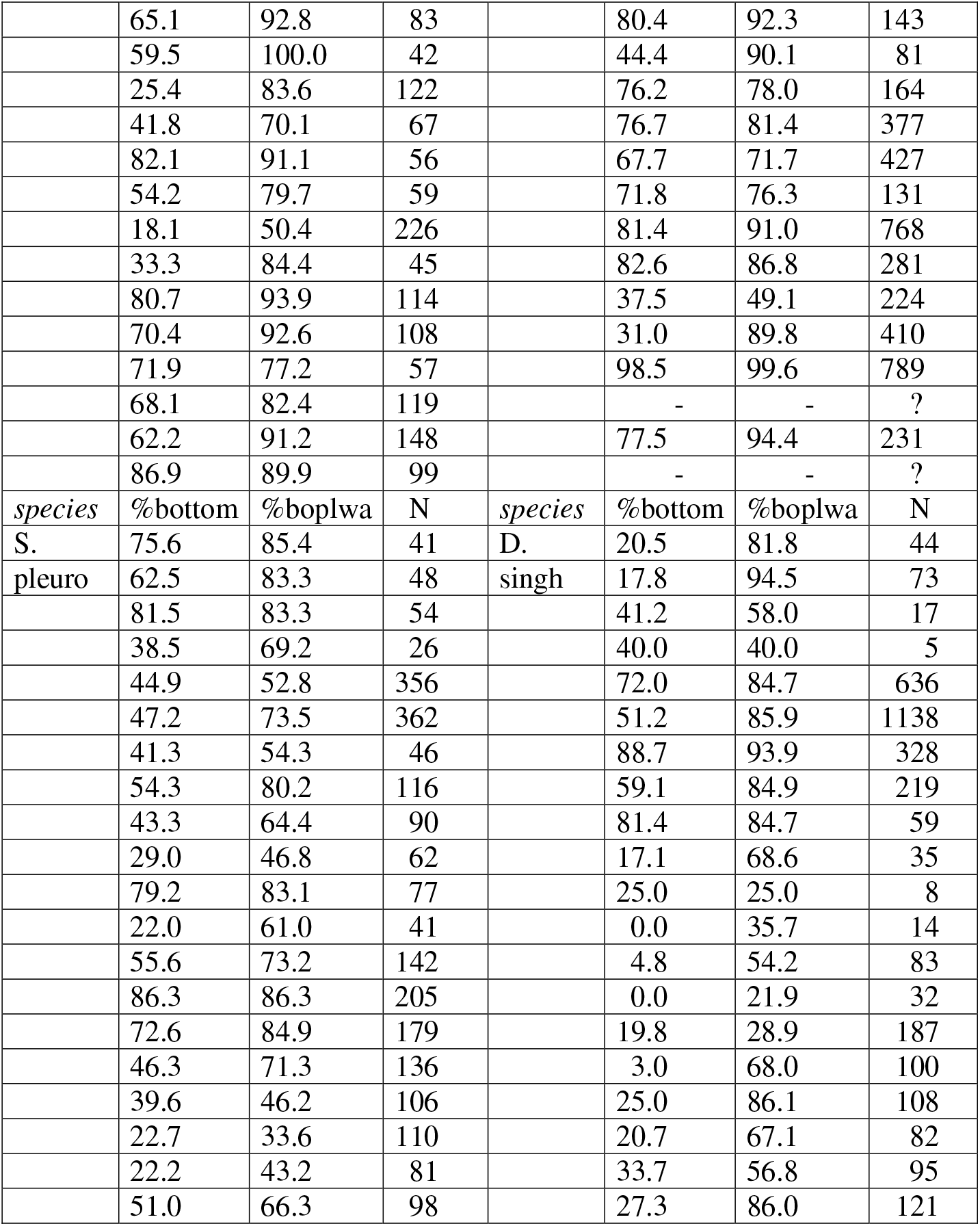
Percentages of snaps directed at bottom and bottom+plants+walls (boplwa) respectively, for 20 samples of all 10 species. P.kama = *Puntius kamalika*; P.chola = *P*.*chola*; P.bimac = *P*.*bimaculatus*; P.dors = *P*.*dorsalis*; P.vitt = *P*.*vittatus*; P.titt = *P*.*titteya*; Pe.nigro = *Pethia nigrofasciata*; Pe.reval = *Pethia reval*; S.pleuro = *Systomus pleurotaenia*; D.singh = *Dawkinsia singhala*.

The percentage ‘free’ follows from the latter by subtraction from 100%. For a graphical representation of the differences between the species, we chose the statistic R from the Friedman statistical test for *k* related samples (figs 1a and b). According to the test, the overall differences between the species are highly significant (*p* < 0.001) for both parameters.

**Fig 1a.**
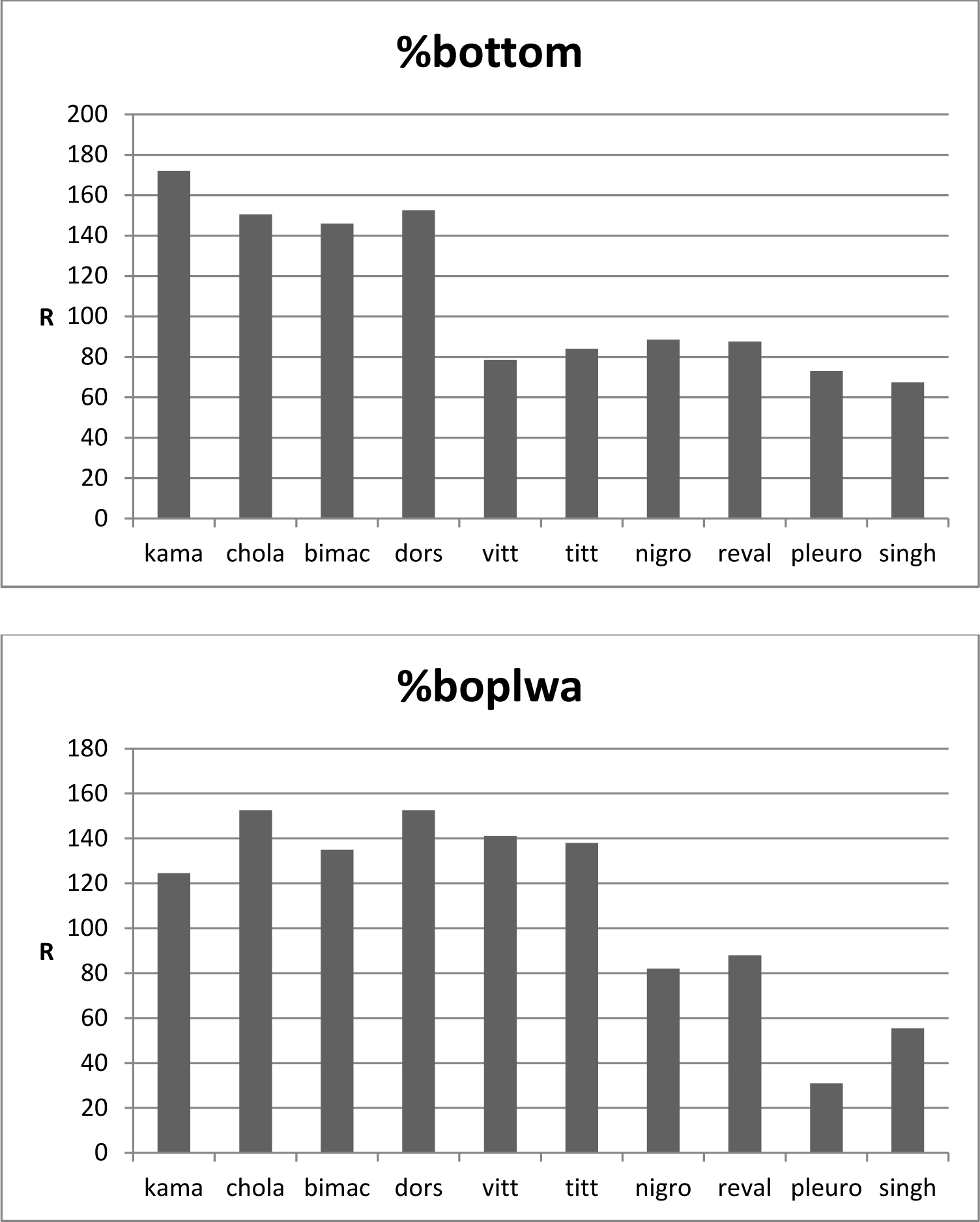
Percentage of snaps directed at bottom over all 10 species. Ordinate: rank statistic R from Friedman two-way analyses of variance over 20 samples per species **Fig. 1b** Percentage of snaps directed at bottom+plants+walls (boplwa) over all 10 species. Rest as Fig. 1a

Both graphs divide the 10 species into at least two groupsof 4 and 6 members, but in a different manner. Specifically, *Puntius vittatus* amd *P. titteya* are on a par with *Pethia nigrofasciata, Pethia reval, Systomus pleurotaenia* and *Dawkinsia singhala* as to % bottom alone, but as for all fixed substrates they join the bottom feeders *Puntius kamalika, P. chola, P. bimaculatus* and *P. dorsalis*. In other words, *Puntius vittatus* and *P. titteya* feed almost exclusively from fixed surfaces, but they are not at all bound to bottom alone. Also, in the % bottom+plant+wall, *P. vittatus* and *P. titteya* differ from both *Pethia* species, *nigrofasciata* and *reval* as well as from *S. pleurotaenia* and *D. singhala*, all of which are *not* confined to fixed substrates. The two *Pethia* species differ from *S. pleurotaenia* and *D. singhala* in that the latter two feed even more from free substrates. Two by two testing of all 10 species with the Wilcoxon matched pairs test confirmed, with few exceptions, all the differences mentioned (tables 2a and b for values of *p*)^7^. Note that the members of the pairs: *vittatus-titteya, nigrofasciata-reval* and *pleurotaenia-singhala* do not differ significantly among each other.

**Table 2a.**
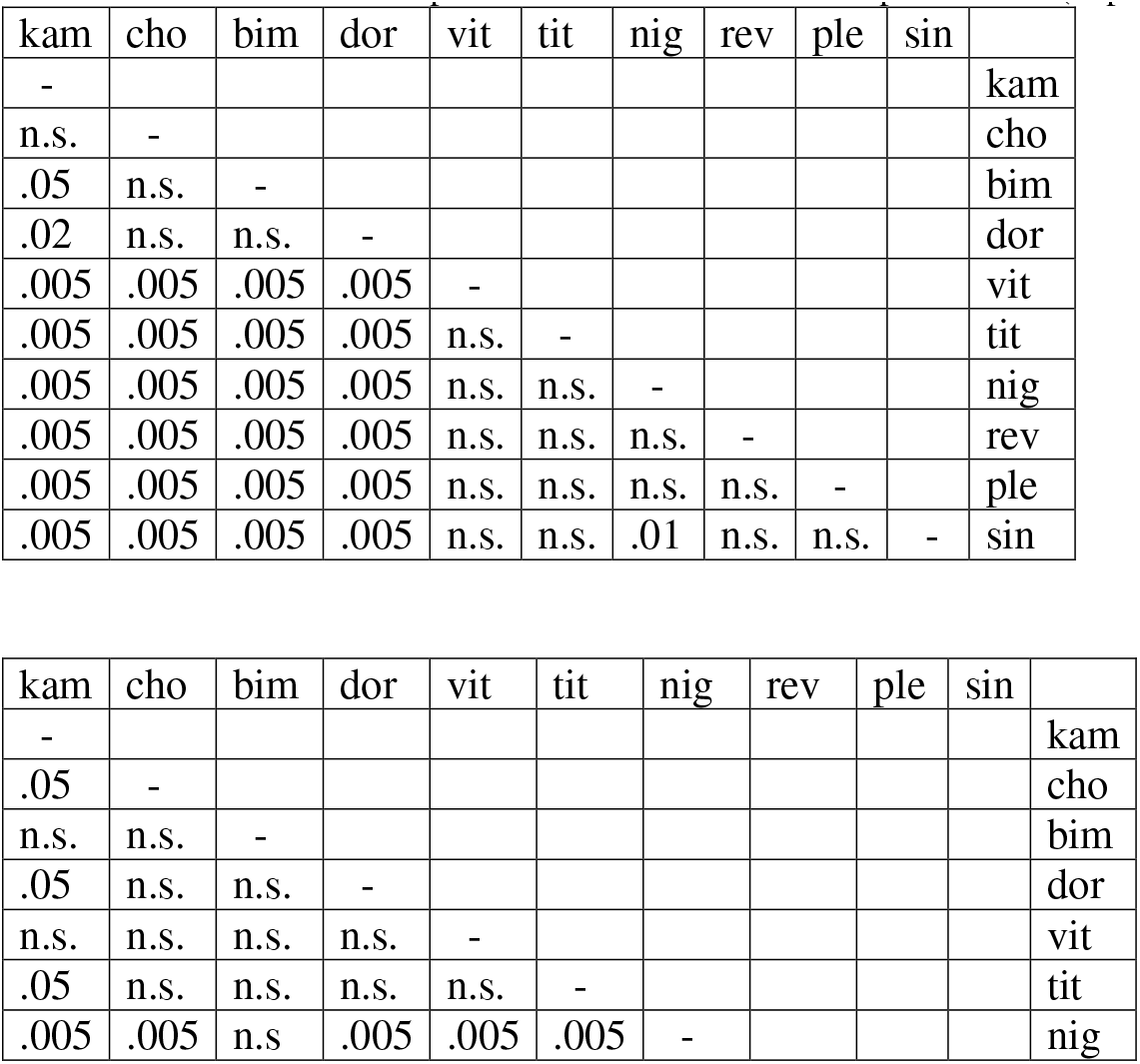
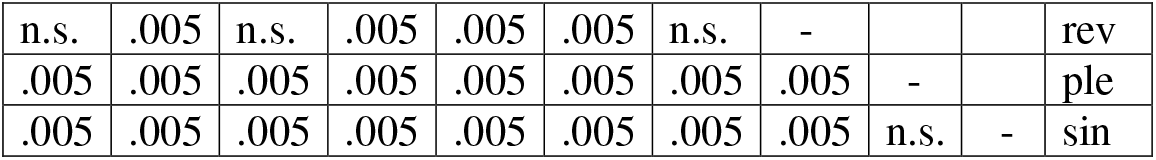
Values of *p* for differences between all species taken two by two over 20 samples. n.s. = not significant. Comparison of values of %bottom. Specific names abbreviated to first three letters. Table 2b. As Table 2a. Comparison of values of %bottom+plants+wall (boplwa).

In summary, we may distinguish four feeding types: a. The typically bottom feeding *Puntius* species *kamalika, chola, bimaculatus* and *dorsalis*, b. *Puntius vittatus* and *P. titteya*, restricted to fixed substrates, but not to bottom alone, c. *Pethia nigrofasciata* and *Pethia reval* doing a considerable proportion of ‘free’ snapping, and d. *S. pleurotaenia* and *D. singhala* which are most free of all from the fixed substrates.

### 2. Other parameters

Besides substrate choice, the 10 species differ among each other in how they search for food, how deep they delve into the bottom, the special moves that assist delving and the processing of food after the snap.

#### Search

Typically, a target is fixated from nearby; when performed at the bottom, this is accompanied by a roughly 30^°^ head-down position (p.00). Table 3 (two samples available) shows the frequencies of searching episodes without snapping for all species. The most surprising is that the four bottom feeding species do very little searching of this sort. More typically, they dip down at the bottom from a normal swimming position (unexpectedly for us), take a mouthful and return the non-digestible matter through the mouth or the gills. They may perform snaps in series, but without searching in between. Possibly, they locate potential food items by sight from a more wide-screen view; or alternatively, they may hunt by olfactory cues.

**Table 3.**
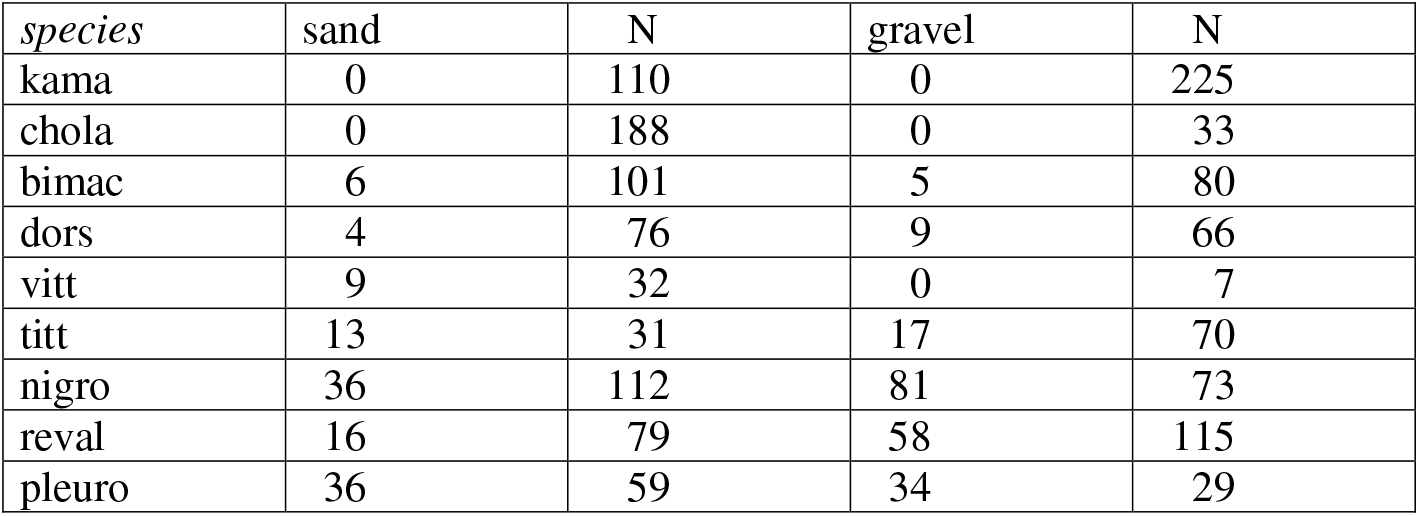
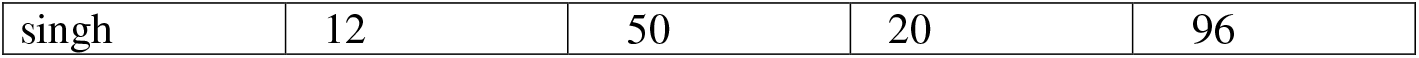
Percentages of episodes that consisted of searching posture without snaps over 10 species. Two samples with sand (left) and gravel (right) bottoms respectively. Abbreviations as in Table 1.

| *species* | sand | N | gravel | N |
| --- | --- | --- | --- | --- |
| kama | 0 | 110 | 0 | 225 |
| chola | 0 | 188 | 0 | 33 |
| bimac | 6 | 101 | 5 | 80 |
| dors | 4 | 76 | 9 | 66 |
| vitt | 9 | 32 | 0 | 7 |
| titt | 13 | 31 | 17 | 70 |
| nigro | 36 | 112 | 81 | 73 |
| reval | 16 | 79 | 58 | 115 |
| pleuro | 36 | 59 | 34 | 29 |
| singh | 12 | 50 | 20 | 96 |

#### Depth

Depth of bottom snaps may vary from S_0_, at the substrate’s surface, to S_5_, down to the gill opening. Table 4 gives the percentages of snaps that exceed S_0_. The two samples available (left: sand; right: gravel) agree in that the highest scores are for three of the four bottom feeders. The exception is *Puntius bimaculatus*. A specific snapping variant in this species that we have observed but was not included in this study, is to let the head drop on the bottom, thus whirling up some debris, from which it may snap some. This may have occurred and been registered as S_0_. Also conspicuous in table 4 are the high scores for *S. pleurotaenia* and *D. singhala* in the sample with gravel. These two species had with distance the largest body size of all, and we presume that for them the gravel was more easy to handle than the sand, since the particle size was not a problem for them, and the rounded shape of the same reduced the cohesion of the medium.

**Table 4.** Percentages of bottom snaps with depth more than zero (%>S_0_) over 10 species in two samples with sand (columns 2-4) and gravel (columns 5-7) bottoms respectively. Rel.shift = difference in positions of all species between orders of standard length and of depth (number of steps forward or back).

| species | $\%>S_0$ | N | rel.shift | $\%>S_0$ | N | rel.shift |
| --- | --- | --- | --- | --- | --- | --- |
| kama | 96.0 | 397 | 4 | 98.0 | 1031 | 3 |
| chola | 95.6 | 800 | 2 | 99.2 | 128 | 3 |
| bimac | 42.6 | 627 | 1 | 38.1 | 404 | 1 |
| dors | 94.3 | 646 | 0 | 92.2 | 258 | 0 |
| vitt | 34.3 | 73 | -1 | 2.1 | 283 | -2 |
| titt | 35.3 | 411 | 1 | 30.3 | 716 | 1 |
| nigro | 47.4 | 310 | 1 | 32.0 | 75 | -1 |
| reval | 18.7 | 171 | 0 | 3.0 | 231 | 1 |
| pleuro | 56.6 | 159 | -3 | 79.4 | 165 | -3 |
| singh | 36.2 | 458 | -5 | 73.2 | 583 | -3 |

This takes us to a comparison between digging performance and specific size. The standard lengths (SL) of the fish were measured (a representative one for each species). Figs 2a and b compare SLs with the digging depths of the species. The general trend is for depth to increase with body size, but *S. pleurotaenia* and *D. singhala* delve less deeply than expected on this trend^8^. Thus, the relatively deep digging of these two species according to table 4 is due to their relatively large size, not to their investing extra effort. Rather, they moderate their effort and invest in variation of substrates. Columns 4 and 7 of table 4 illustrate the discrepancies between the parameters depth and SL. Apparently, the smaller of the four bottom feeding species dig deeper than expected on the basis of their body sizes (positive figures). Conversely, *S. pleurotaenia* and *D. singhala* dig less deep (negative).

**Fig 2a.**
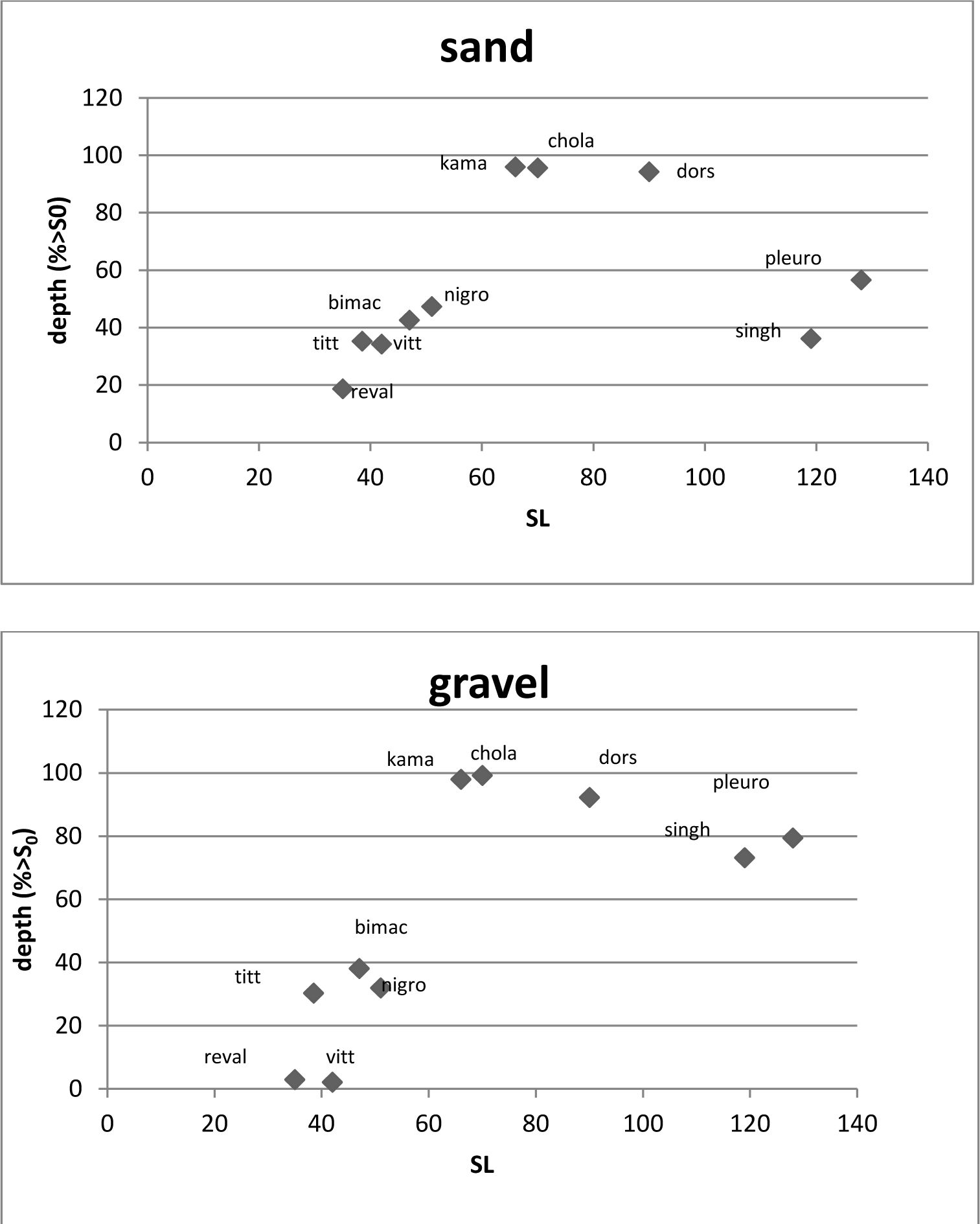
Sand bottoms. Standard Lengths (SL) of fish and the depths of snaps (percentage of snaps with depths more than zero: %>S_0_) on abscissa and ordinate respectively **Fig.2b** Gravel bottoms. Standard Lengths (SL) of fish and the depths of snaps (percentage of snaps with depths more than zero: %>S_0_) on abscissa and ordinate respectively

#### Digging techniques

The most common movements assisting digging were Butting (B), Head-shaking (H) and Pushing (Pu)^9^, (BHP). (For Sieving (Si) see below). The percentage of snaps accompanied by one or more of them is in table 5 (left: sand; right: gravel) for the same two samples as in table 4. Only with the gravel bottom, the *Puntius* species *kamalika, chola* and *bimaculatus* perform considerable percentages of BHP; especially *kamalika* and *chola* do less of it in the sample with sand.

**Table 5.** Percentages of bottom snaps with Butting, Headshaking and/or Pushing (%BHP) and with Tugging and/or Wrenching (T+W) over all species, in two samples over sand (columns 2-4) and gravel (5-7) bottoms respectively.

| *species* | %BHP | %T+W | N | %BHP | %T+W | N |
| --- | --- | --- | --- | --- | --- | --- |
| kama | 6.8 | 0.0 | 397 | 24.6 | 0.2 | 1031 |
| chola | 8.6 | 0.0 | 800 | 35.9 | 0.0 | 128 |
| bimac | 26.5 | 1.6 | 627 | 34.4 | 0.2 | 404 |
| dors | 1.1 | 0.8 | 646 | 3.1 | 0.4 | 258 |
| vitt | 19.8 | 2.5 | 81 | 6.7 | 3.2 | 283 |
| titt | 1.7 | 0.7 | 411 | 3.2 | 1.8 | 716 |
| nigro | 1.3 | 36.5 | 310 | 18.7 | 5.3 | 75 |
| reval | 0.6 | 4.7 | 171 | 3.0 | 20.3 | 231 |
| pleuro | 7.5 | 1.3 | 159 | 10.3 | 0.0 | 165 |
| singh | 1.3 | 3.5 | 458 | 1.9 | 3.9 | 583 |

It may be noted that *Puntius dorsalis* hardly performs any BHP in either sample (though some was combined with sieving). Instead, it Sieves (Si), sucking in matter from the substrate and spilling it through the gill openings, filtering out edible particles with the gill rakers (41.1% and 62.1% of snaps with sand and gravel respectively). None of the other species does this to any appreciable extent (3.5% in *P. chola*, rest under 1% if any). Another technique is tugging (T) and wrenching (W) at plants (table 5). It is a specialty of the two *Pethia* species *nigrofasciata* and *reval*. These tug in particular at short stems or roots sticking out of the bottom at the base of macrophytes.

The final visible phase of eating is the processing of the snapped-up matter in the mouth. Table 6 shows percentages of occurrence of chewing (C), spitting (Sp) of bottom material and sieving (Si) for the same two samples as before. They are expressed as percentages of snaps that are followed by any or combinations of these. In *P. dorsalis* 73.7% and 51% of snaps on sand and gravel respectively are accompanied by Si, singly or in combination with other techniques. Of the other species only *P. chola* scored 1.5%. Comparison with table 4 (depths of bottom snaps) suggests a concordance between the two parameters: deeper diggers tend to process their food by more C, Sp and/or Si (figs. 3a and b). The correlation is significant (Spearman *r*_*S*_ = 0.973 resp. 0.842; *p* < 0.01 in both cases). (Note that the figures for *P. dorsalis* include sieving, and the percentages of Si may include combinations with C and/or Sp). *P. dorsalis* is the only species practicing Si to a considerable extent both in delving and the processing of food.

**Fig 3a.**
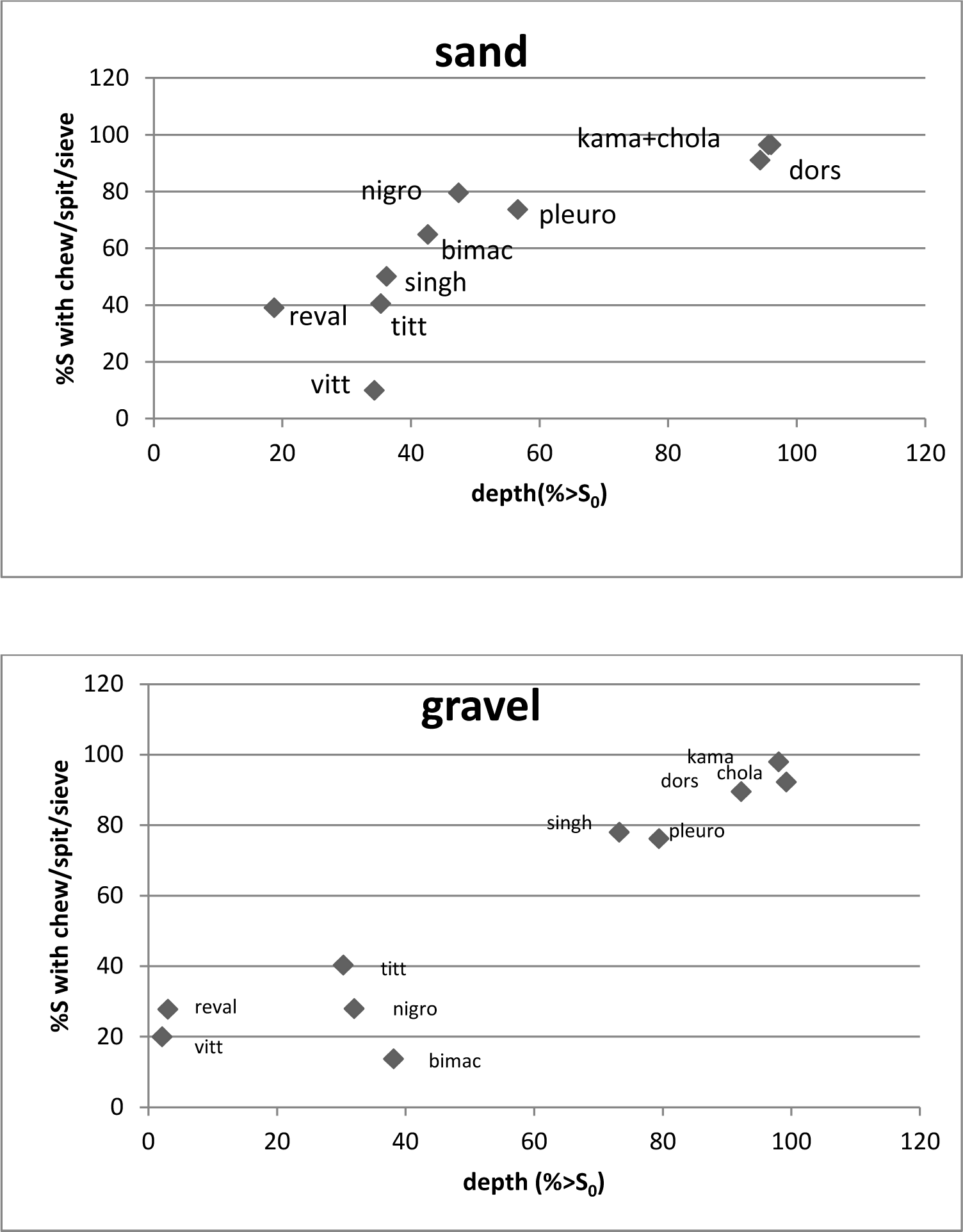
Sand bottoms. Correlation between depths of delving (table 4) and frequencies of food processing movements (Chewing, Spitting and/or Sieving)(table 6) **Fig. 3b** Gravel bottoms. Correlation between depths of delving (table 4) and frequencies of food processing movements (Chewing, Spitting and/or Sieving)(table 6)

**Table 6.** Percentages of bottom snaps followed by Chewing, Spitting, Sieving, or combinations thereof (%C/Sp/Si) over all species. Two samples with sand (columns 2 and 3) and gravel (columns 4 and 5) bottoms respectively. Data from different parts of observation sessions relative to percentages depth (table 4); hence different Ns.

| *species* | %C/Sp/Si | N | %C/Sp/Si | N |
| --- | --- | --- | --- | --- |
| kama | 96.5 | 173 | 97.9 | 144 |
| chola | 96.5 | 462 | 92.2 | 58 |
| bimac | 64.9 | 138 | 13.7 | 190 |
| dors | 91.1 | 190 | 89.5 | 229 |
| vitt | 10.0 | 10 | 20.0 | 15 |
| titt | 40.6 | 187 | 40.3 | 325 |
| nigro | 79.6 | 54 | 28.0 | 25 |
| reval | 39.1 | 115 | 27.8 | 36 |
| pleuro | 73.7 | 19 | 76.2 | 63 |
| singh | 50.2 | 291 | 78.0 | 129 |

Correlations with and between the other parameters (%BHP+Si for incidence of delving techniques, and %+C/Sp/Si for food processing) are less clear or absent. %BHP+Si *vs* %>S_0_ is not significant on sand bottoms, and just significant on gravel (*r*_S_ = 0.636; *p* < 0,05). Correlations between %BHP+Si and %+C/Sp/Si are not significant.

The effects of BHP and Si on depth can also be investigated by comparing, for each species separately, the depths of snaps with and without them. χ^2^ tests were done, for each species and for the two samples available, on the difference made by B and/or H^10^, and also by Si in *Puntius dorsalis*, 22 cases in all. The majority (18) were significant at *p* < 0.001 in the expected direction. Only those of *Pethia nigrofasciata* and *P, reval* were not significant (for lack of sufficient data), and one case of *Dawkinsia singhala* gave a (significant) reverse effect.

In summary, the other parameters, in combination with the substrate preferences define feeding behavior profiles of each of the 10 species. The similarity in substrate choice between the four predominantly bottom feeding *Puntius* species is corroborated by the relative absence of close optical searching, the presence of relatively deep delving and much chewing and spitting after the snap, the two last items with the exception of *P. bimaculatus* That species does perform digging movements (BHP) in considerable frequency, but these result in comparatively shallow snaps. The cause may well be in the morphology of its mouth, but we have no data on this. *P. dorsalis* differs from the other three through the predominance of sieving and filtering. It has the wide, broad-based mouth cavity that optimizes sucking. The two *Pethia* species *nigrofasciata* and *reval* agree in the frequent occurrence of tugging and wrenching at the base of rooted vegetation. *P. nigrofasciata* does this more often than *P. reval*. In general, the former of the two delves a little deeper than the other and chews/spits more, often with some delay between snapping and spitting.

## Discussion

The fact that species can be arranged in feeding types raises the question of coexistence of species of the same type in nature. Co-occurrence and species associations in the hill country of Sri Lanka were studied by Schut *et al*. (1984). A similar exercise on a smaller scale for the lowland species was offered by Kortmulder & Robbers (2011).

Are species of the same feeding type associated or separated in nature? *Puntius titteya* and *P. vittatus*, to begin with, inhabit widely separated waters, the former in the peaty hill marshes far above flood level, the latter everywhere in the lowlands. In fact, according to Schut *et al*., *P. vittatus* correlates negatively with all hill stream barb species. The correlation between *Pethia nigrofasciata* and *P. reval* is indifferent, meaning that they do meet at some places, but their typical haunts -respectively winding hill streams shaded by trees well above flood level and the mouths of tributaries near flood level -are separate. *Systomus pleurotaenia* associates with *Pethia nigrofasciata* and *Puntius dorsalis* in the hill streams already mentioned, and is separate from *Dawkinsia singhala*, which associates with *Pethia reval* or *P. cumingii* further downstream.

Of the four bottom feeding *Puntius* species *kamalika, chola, bimaculatus*, and *dorsalis* the first two are typically lowland species. Kortmulder & Robbers (2011) suggest that *P. chola* prefers clear waters of all sizes, while *P. kamalika* is restricted to turbid parts of large rivers. Juvenile and young adult *P. bimaculatus* share the high hill marshes with *Puntius titteya*, but the adults are found in fast, stony brooks connected to the same marshes. A little lower, in somewhat more quiet streams, they may also be found together with *Puntius dorsalis* (personal observations by S.S De Silva and the first author; see also Kottelat & Pethiyagoda (1991: 302) where *Pethia bandula* mingles with the same two species). *Puntius dorsalis*, finally, occurs in a relatively wide range of habitats and associations: from the just mentioned combination with *P. bimaculatus*, through the winding hill stream association to the lowlands as far as the water is deep and clear (Kortmulder & Robbers, 2011). We suggest that it is the sucking and sieving method of feeding that enables *P. dorsalis* to coexist with *P. bimaculatus* and share in other associations without much competition. Through its adaptation, unique among the species concerned, it delves as deep as any of the others and certainly deeper than *P. bimaculatus*, which is the most superficial delver among the bottom feeders.

Comparing between species, the only clear correlation between feeding movements is the positive one between depth of digging and processing by chewing/spitting/sieving. The relationship is readily interpretable and underlines the similarities between the bottom-preferring species. Butting, Head-shaking, Pushing and Sieving serve to dig deeper. That the correlation between these moves and digging depth when comparing species does not confirm this (p. 00) may be due to other factors influencing depth, such as body size and effort.

We conclude that all species that have similar modes of feeding, are largely separated as to their respective habitats. This conclusion tallies well with what others have found: co-existing species differ in their feeding behaviour (Moyle & Senanayake 1984; Schut et al. 1984), but it is arrived at in an independent manner and on the basis of data of a different sort.

Table 7 summarises the diets as collected from published data. In most cases the intestinal remains reflect what one would expect on the basis of feeding behaviors in the aquarium. For instance, the preference of *P. dorsalis* for chironomid larvae among animal preys may well result from the depth of its delving, because these larvae live below the bottom surface. Ephemeropteran and trichopteran larvae, in contrast, live at the bottom surface, the latter typically nestling against rooted plants (J. Vijverberg, pers. comm.). The same may be true for the preference that *Pethia nigrofasciata* showed for chironomid larvae, contrasting with *P. cumingii*, which preferred ephemeropterans (Vijverberg *et al*., 2019). The tugging and wrenching techniques used by both species may serve the detection of trichopteran larvae, but this does not show in the gut analyses, possibly because trichopterans were generally scarce in the pools studied, where both ephemeropterans and chironomids were abundant (Vijverberg, pers. comm.). As to the other species, in addition to some surprises (*P. dorsalis, P. titteya*), there are many details that could not be predicted from the aquarium-observed behavior, but depend on what the environment had to offer. Apparently, our aquarium studies cannot *replace* field work. Analyses of gut contents of fish in their natural setting remains necessary, as well as the here recommended sampling of environmental resources and the observation of fish as they feed in nature.

**Table 7.**
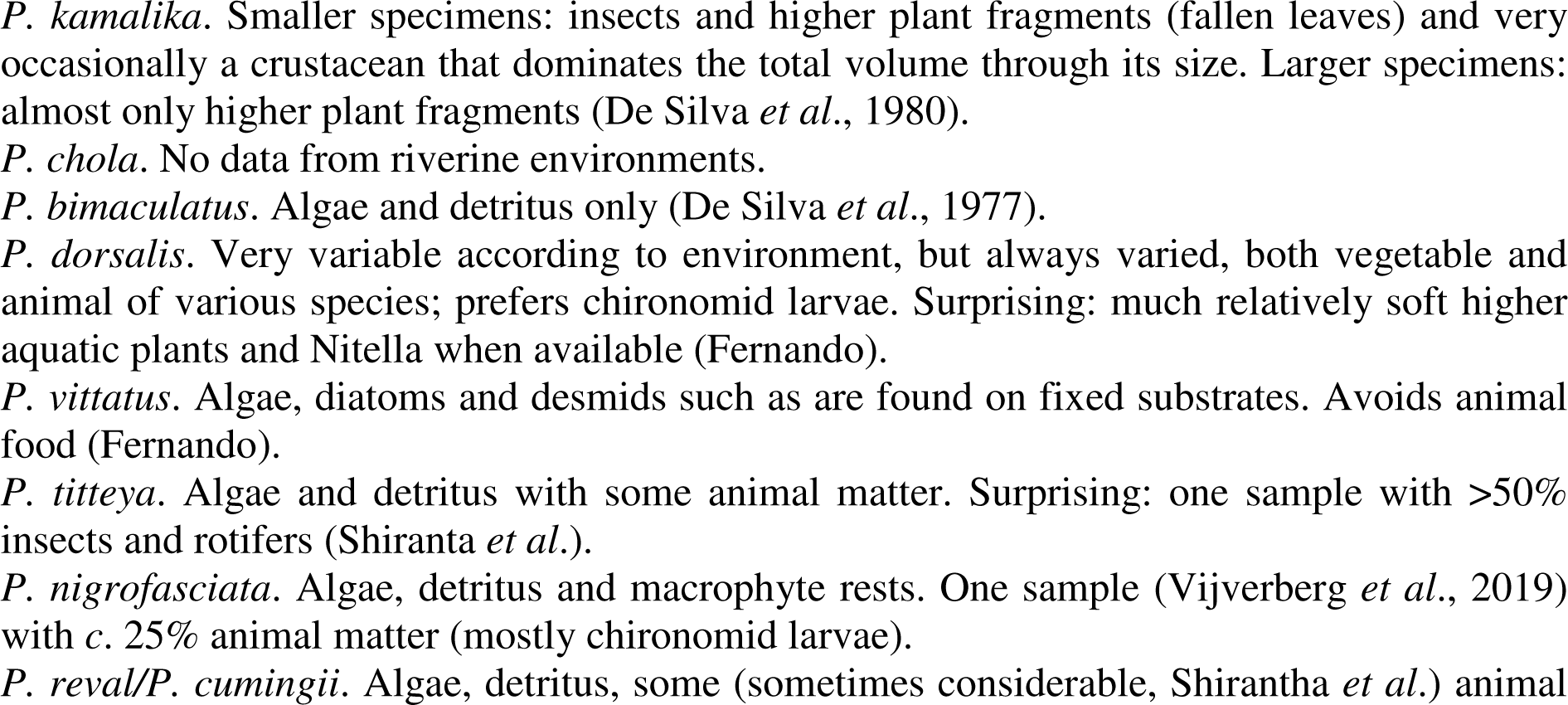
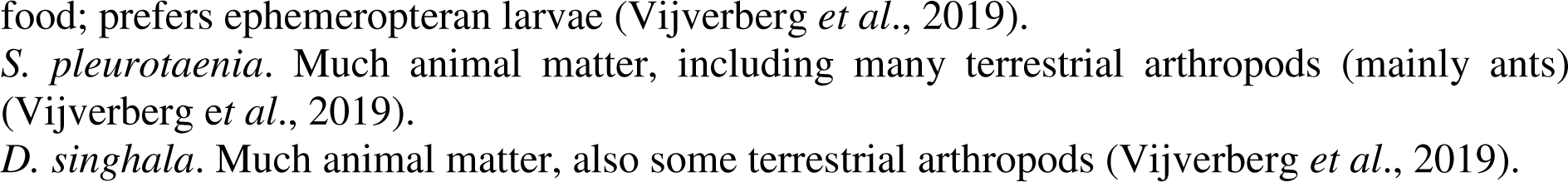
Data on natural diets established by gut analyses, collected from the literature.

### What then is the use of aquarium observations?

First, the behavior can be observed in much more detail in aquaria, and observing the different species under similar conditions in the lab brings out the natural inclinations of each species, so that one may be sure they are evolutionary adaptations. The same is true for morphology, but behavior is nearer to the process of food intake.

Identification of food remains in the gut does not tell from where they were taken. The presence of sand or fallen leaf fragments may give further indications, but observing the behavior of living fish is decisive. This is an important point in the estimation of dietary overlaps between co-occurring species. Fish that take prey of the same species but from different substrates or depths can hardly be considered as competing, at least not in the current generation of the prey species.

Finally, information on feeding behavior is one step nearer to a complete understanding of the entire feeding process. What remains to be studied is the phase where a fish may actually choose between two or more items that are equally accessible. Also this will be best studied under controlled aquarium conditions.

## Acknowledgements

The financial support of the ‘Dr. J. L. Dobberke Stichting voor Vergelijkende Psychologie’ is gratefully acknowledged. Thanks are due to Dr. Jacobus Vijverberg for useful communication throughout the preparation of this paper; to Mr. Yuri Robbers who advised as to statistical testing; to both for reading the ms in draft and commenting on its content. The assistance of Mr. Jalb Schut in the observations is acknowledged with thanks.

## Declarations

### Authorship statement

Both authors contributed to the study conception and design. Material preparation, data collection and primary analysis were performed by Peter van der Wiele. The first draft of the manuscript was written by Koenraad Kortmulder, and the second author commented on a previous version of the manuscript. Both authors agree on the final manuscript.

### Funding and/or Competing Interests declaration

The authors have no financial or proprietory interests in any material discussed in this article. Partial financial support (a salary for the second author) was received from the Dr. J. L. Dobberke Stichting, a Dutch non-profit fund supporting ethological studies.

### Ethics Approval declaration

The experiments on which this study is based were performed in the early nineteen-eighties, when there was no external ethical screening of our institute. However, the fish used in this study were kept and tested under near-natural conditions and we think no ethical approval is required.

### Data Availability Statement

The data on which this article is based were first laid down in two voluminous reports type-written in Dutch in 1982 and 1983 respectively. This was the usual format in our institute at the time. The reports are in the possession of the first author, who is willing to have them checked.

However, there are no data with any bearing upon the results of this article, generated and analysed during the study, that are not included in the published paper.

## Footnotes

2 Then all were called *Puntius*; only *Puntius sarana* (now *Systomus sarana*) and *Puntius thermalis*, both lowland species, we did not have. New since then are *Pethia bandula*, which is closely related to *Pethia nigrofasciata*; *Dawkinsia srilankensis, Systomus asoka* and *Systomus martenstyni*, all three living in fast-flowing, deep waters in hill country, possibly vicarating with *Systomus pleurotaenia* or *Dawkinsia singhala* (then: *Puntius filamentosus*); and *Puntius kelumi* Pethiyagoda et al., 2008, similar to *P. dorsalis. Puntius kamalika* has been distinguished from *P. amphibius* from India by Silva et al., 2008. *Pethia reval* has been split off from *Pethia cumingii* by Meegaskumbura et al., 2008; the two have comparable habitats in different river systems.

3 Twice a day, 15 minutes before the first recordings and after the last one of the day.

4 New specimens of this species had been imported along with the others, but we are not sure whether these or the lab stocks were used, probably the latter.

5 The lab stock were *P. reval*, with red fins. *Pethia reval* and *P. cumingii* are very closely related. In the lab, they hybridize easily and produce viable males and females, in contrast to other *Pethia* species of the region (Kortmulder, 1972). They have similar ecological roles in their respective river systems (De Silva *et al*., 1985; Schut *et al*., 1984).

6 In one tank (part I, first series of *Puntius chola*) the macrophytes were overgrown by the algae; for the second series the latter were largely removed and new *Vallisneria* planted.

7 The few exceptions *e*.*g. kamalika vs chola, dorsalis* and *titteya*) are significant at a much lower level.

8 or the bottom-feeding species deeper, but that is not the point here.

9 Butting is a straightforward move at or into the substrate, energized by the tail, Head-shaking an oscillating, sideways movement, with the mouth touching the substrate. Pushing shoves some sand forward creating a small pit below the fish’s mouth.

10 Pushing (Pu) occurred in only two species, *Puntius chola* and *Systomus pleurotaenia*, and was rare even there. It was therefore omitted from this statistical procedure.

